# Population genomics of the rice landrace Acuce reveals exceptional dynamics of immune receptors

**DOI:** 10.1101/2024.02.15.579491

**Authors:** Stéphane De Mita, Shengchang Duan, Xiang Li, Rémi Pélissier, Laura Mathieu, Jingyu Li, Linna Ma, Zihang Li, Jingjing Liao, Xiahong He, Youyong Zhu, Chengyun Li, Yang Dong, Jean-Benoit Morel, Huichuan Huang

## Abstract

In cultivated plants such as *Oryza sativa*, landraces harbour high levels of diversity, in sharp contrast with genetically uniform modern varieties. The Yuanyang terraces (China) are renowned for traditional farming involving landraces and low disease incidence. To understand the molecular basis of these resistance levels, we explored the genomic diversity of Acuce, one of the oldest landraces in the region. Based on whole-genome resequencing of over 200 Acuce plants, we evaluated the diversity of immunity and agronomical genes with respect to the rest of the genome. In parallel, we analysed the genetic basis of pathogen resistance and agronomical traits. Acuce exhibits high levels of diversity, reaching 38% of the worldwide levels. We uncovered an excess of diversity and signatures of directional selection associated with immunity genes, in contrast with agricultural traits. Furthermore, cultures of mixed genotypes exhibited superior performances for both pathogen resistance and agricultural traits. We showed that, within one single landrace, traditional farmer practices selected optimal agricultural traits while maintaining diversity enabling pathogen resistance. This study elucidates the complex interplay between genetic diversity, natural selection related to pathogens, and selection for agricultural traits within traditional landraces, providing insights into sustainable agricultural practices in diverse cropping systems.

---

Keywords: landrace, diversity, immune receptor, pathogen resistance, *Oryza sativa*

## Introduction

Landraces represent a traditional way of maintaining diversity in cultivated systems (<u>McCouch et al., 2020</u>). While they do not result from deliberate improvement, landraces are both locally adapted and genetically diverse, but are still identified and unambiguously delineated by farmers (<u>Harlan, 1975</u>; <u>Villa et al., 2005</u>). Local or regional landraces of rice (*Oryza sativa*) have been analysed to some extent in Borneo and Madagascar islands (<u>Mather</u> <u>et al., 2010</u>; <u>Thomson et al., 2009</u>). However, there is only limited knowledge on how diversity is structured within individual landraces, and none at the genomic scale (<u>Pusadee et</u> <u>al., 2009</u>). Specifically, it is not known whether individual landraces are homogeneous or, instead, harbour substructure with internally differentiated groups. Moreover, the molecular functions that are genetically diversified in landraces are not known. Furthermore, the question of whether landraces can be engineered in modern selection programs has not been addressed to date.

The Yuanyang terraces (Yunnan Province, China, hereafter YYT) were shaped by the agricultural practices of the Hani ethnic group (<u>Wang, 1999</u>). They were classified as a “cultural landcape” by UNESCO in 2013 and are known for being home to a wide range of rice landraces (<u>Jiao et al., 2012</u>). Quite importantly, the available records indicate that the YYT only suffers 1% losses caused by disease (<u>Li, 1990</u>), dramatically less than the 32% losses at the world level (<u>Savary et al., 2019</u>). High genetic diversity of some YYT landraces has been documented in previous reports using a limited number of molecular markers (<u>Gao</u> <u>et al., 2012</u>; <u>Zhang et al., 2007</u>), along with signatures of substructure. Recently, an analysis combining RenSeq targetting genes involved in immunity and genotyping-by-sequencing, has shown extensive levels variation and signatures of balancing selection in immunity genes within four landraces cultivated in the YYT and belonging to the XI group (standing for Xian / Indica) of *Oryza sativa* (<u>Gladieux et al., 2024</u>). Among them, Acuce (literally “God rice” in the Hani language and “Moon rice” in Mandarin) is a prevalent landrace in YYT (<u>Ali</u> <u>et al., 2023</u>; <u>Alonso et al., 2019</u>). It has been grown in YYT for more than 130 years (Figure S1A) and appears to have remained immune to the major fungal pathogen *Pyricularia oryzae* (<u>Ali et al., 2023</u>). Although cultural practices in YYT can be hypothesized to provide an efficient disease control, it is not clear whether this effect is due to a particular content of genes mediating immunity. A genome-wide survey of nucleotide diversity is required to analyse the structure of diversity within landraces and the effect of disease-mediated selective forces on genes involved in immunity. Within the YYT, the genomic diversity is analysed across 91 landraces by <u>Huang et al. (2025)</u> and here we focus on Acuce to provide a detailed survey of diversity occurring within a single landrace. The Acuce landrace offers a unique opportunity to understand how the definition of a landrace by farmers is consistent with the structuring of genomic diversity, the molecular basis of stable and robust pathogen resistance, and the phenotypic and genotypic traits through which high intra-landrace diversity is expressed.

In this study, we relied on over 200 Acuce plants sampled from various villages of the YYT (Figure S1B) to analyse the structure of genomic diversity based on Illumina genome resequencing. These data were compared with the genome of representatives from other landraces of the YYT and accessions from worldwide collections (Figure S1E). We conducted a detailed analysis of genomic diversity structure, compared regions involved in pathogen resistance with the rest of the genome and analysed footprints of selection. In addition, we provide hints about genotype-phenotype relationship by examining genetic variance of disease resistance traits in comparison to agronomical traits. We put results concerning Acuce in perspective by examining a variety obtained through modern breeding, HongYang 3 (hereafter HY3; Figure S1C), which was introduced in 2014 in the YYT.

## Material and Methods

### Sampling of Acuce, HY3 and other rice genomes

Acuce and HY3 plants were sampled from a small sector (∼8×13 km) in the Yuanyang County (Yunnan, China; Figure S1B). This area is similar to that described by (<u>Jiao et al., 2012</u>). A total of 227 Acuce and 30 HY3 plants were included in the sample. The 227 Acuce samples include 89 single-seed descents that have been propagated for 8 years starting from seeds collected in Qingkou village. This samples are referred to as inbred. All other Acuce and HY3 were collected in 2015 and 2016 (Figure S1D). The landrace or variety status was based on input from local farmers. In order to clarify the organization of the diversity with Acuce with respect to larger groups, we included 252 samples from <u>Huang et al. (2025)</u>: 192 belonging to YYT clusters (including 4 Acuce samples) and 60 genomes representative of rice world diversity (50 from the XI group and 10 from the GJ group, the latter standing for Geng / Japonica) (<u>Wang et al., 2018</u>). Overall, 509 genomes were included in the analysis (Table S1), and *Oryza barthii* (accession number: 65489_Ob_516W1) (<u>Cubry et al. , 2018</u>) was used as an outgroup.

### Field phenotyping

The 89 inbred Acuce lines were sown and transplanted as pure stands in three separate fields in Qingkou village in 2022 (Figure S3A). Plant height, tiller number, and thousand-seed weight were measured in the fields. Four indicators associated with micro-organism associated diseases (blast, neck blast, brown spot, and smut) were surveyed using a grading scale ranging from level 0 to level 5 during the field survey. Subsequently, we converted the disease grades to disease indices by adopting the method from <u>Chiang et al. (2017)</u>. Six Acuce inbred lines were selected (A15, A36, A49, A69, A74, and A87) and assembled into 6 binary mixtures. They were evaluated for three years. For each mixture, one line was considered as focal and was scored for phenotype, the other line(s) of the mixture being treated as neighbours. For each focal line and year, the number of plants scored for establishing phenotypes ranged from 195 to 441. We used linear modelling in R to scrutinize the impact of year, field, pure/mix, and focal plant on the measured metrics using the model equation: log(1+value) ∼ year + field + condition × focal. Subsequently, the R package lsmeans was utilized to compute the least squares means for plotting. The p-values were extracted using lsmeans and the pairs function in R. The R package FactoMineR (Version: 2.9) (<u>Lê et al.,</u> <u>2008</u>) was employed for PCA analysis with default parameters. For correlation calculations using Pearson’s correlation coefficient, we utilized the R package WGCNA (Version: 1.72.1) (<u>Langfelder and Horvath, 2008</u>). The *t*-test was applied to examine the significance of differences in data adhering to a normal distribution, while the Wilcoxon test was employed for the remaining non-normally distributed data.

### DNA sequencing and mapping on reference genome

Young leaves were collected from plants and snap-frozen in liquid nitrogen. Total DNA was extracted uisng the DNAsecure plant kit (TIANGEN, Beijing). For each accession, 2 µg of genomic DNA were used to construct a sequencing library following the manufacturer’s instructions using the NEBNext Ultra DNA Library Prep Kit (NEB Inc., USA). Paired-end sequencing libraries with an insert size of approximately 400 bp were sequenced on an Illumina HiSeq 4000 sequencer. The average coverage per genome was 14.5X (Table S11). Paired-end resequencing reads were cleaned using fastp (Version: 0.12.2) (<u>Chen et al., 2018</u>). Cleaned paired-end reads were mapped to the XI rice genome Shuhui498 (alternatively named R498) reference genome (<u>Du et al., 2017</u>) with BWA (Version: 0.7.10-r789) (<u>Li and</u> <u>Durbin, 2009</u>). SAMtools (Version: 1.3.1) (<u>Danecek et al., 2021</u>) was used to convert the mapping results into BAM format. Duplicated reads were marked with the Picard package (picard.sourceforge.net, Version: 2.1.1) after sorting BAM files using the Picard package. After BWA alignment, the reads around indels were realigned, realignment was performed with the Genome Analysis Toolkit (GATK, Version: 3.3-0-g37228af) (<u>McKenna et al., 2010</u>) in two steps. The first step used the RealignerTargetCreator package to identify regions where realignment was needed, and the second step used IndelRealigner to realign the regions identified in the first step, which produced a realigned BAM file for each accession. Variation detection followed the best practice workflow recommended by GATK (<u>McKenna et al.,</u> <u>2010</u>). The SNP variants were called for each accession using GATK HaplotypeCaller. A joint genotyping step for comprehensive variations union was performed on the gVCF files. During the filtering step, the SNP filter expression was set as “QD < 5.0 || MQ < 40.0 || FS > 100.0 || SOR > 3.0 || MQRankSum < -5.0 || ReadPosRankSum < -5.0 || QUAL < 30”. SNPs with none bi-allelic were removed. The SNPs were then cleaned for hidden paralogy as described by <u>Huang et al. (2025)</u>.

### Phylogenetic analysis

The pairwise distance matrix was computed between 510 samples (including *Oryza barthii*) over all SNPs as the proportion of pairwise differences among all SNPs where the given pair of samples have non-missing genotypes. The neighbor-joining tree was computed using the neighbor program of the PHYLIP package (Version: 3.698) (<u>Felsenstein, 1993</u>). *O. barthii* was used as outgroup to root the tree using EggLib (Version: 3.1) (<u>Siol et al., 2022</u>) and the tree representation was realized using the online tool iTol (Version: 5) (<u>Letunic and Bork,</u> <u>2021</u>).

### Structure analysis

Structure analysis was performed using two complementary methods: sNMF (Version: 2.0), as available within the R package LEA (Version: 3.14.0) (<u>Frichot et al., 2014</u>) and Admixture (Version: 1.3) (<u>Alexander et al., 2009</u>). Both programs were used with K ranging from 1 to 15 and 10 replicates. sNMF was run with the parameters: alpha=10, iterations=1000, tolerance=0.000001 and Admixture with cv=10. The analysis was run on 494 samples, excluding the 10 GJ samples, as well as three Acuce (B73, 396, and A3) and two worldwide (304 and B170) samples that appear at a basal position of the phylogenetic tree, indicating a potential taxonomic error. After thinning, 27,522 diallelic sites were included in the analysis. Based on correlations of assignment coefficients, we used *ad hoc* Python code to map clusters with each other over the replicates, the two methods, and the successive values of K to obtain a consistent set of clusters. Individuals were assigned to the cluster with the highest coefficient if it was at least 0.75, or if the gap with the second-best cluster was larger than 0.20.

### Analysis of nucleotide diversity

Only diallelic SNPs were considered in the analysis. Diversity statistics were computed using EggLib (Version: 3.1) (<u>Siol et al., 2022</u>). For each group, we computed the average number of non-missing samples (nseff), number of polymorphic sites (S), total nucleotide diversity (*π*), average unbiased heterozygosity (He), and Tajima’s D (<u>Tajima, 1989</u>), which is an index of bias in the allele frequency spectrum. To measure population differentiation, we computed *F_ST_* as in <u>Weir and Cockerham (1984)</u>, G’_ST (_<u>Hedrick, 2005</u>_)_ and Jost’s D (<u>Jost, 2008</u>), the latter being designed to be less sensitive than the former two to within-population levels of diversity, and both unstandardized (Dxy) and net (Da) pairwise distance (<u>Nei, 1987</u>).

### Acuce genome construction

The Acuce A15 inbred line was used as a reference for PacBio sequencing. A total of 50 mg DNA was used to construct sequencing libraries. Sequencing was conducted on PacBio Sequel platform, and the resulting reads were assembled using MECAT2 (Version: 2019.3.14) (<u>Xiao et al., 2017</u>). The contig was polished using Pilon (Version: 1.21) (<u>Walker et al., 2014</u>) with Illumina reads from the A13 inbred line. For chromosome-scale assembly, Hi-C data were prepared by NowBio Biotechnology Co., Ltd. (Yunnan, China), involving extraction of high molecular weight DNA, digestion with DpnII, biotinylation, dilution, and random ligation of digested fragments. Biotinylated fragments were then enriched and sheared to 300–500 bp for sequencing on Illumina NovaSeq platform. The polished sequences were further scaffolded using Hi-C data based on 3d-dna pipeline (<u>Dudchenko et al., 2017</u>). First, Hi-C reads were mapped to the polished assembly with juicer (Version: 1.5.6) (<u>Durand et al., 2016</u>), and then a candidate chromosome-length assembly was generated automatically using the 3d-dna pipeline (Version: 180922) (<u>Dudchenko et al., 2017</u>). Manual review and refinement of the candidate assembly was performed using Juicebox Assembly Tools (Version: 1.9.1) (<u>Durand et al., 2016</u>) for quality control and interactive correction. The genome assembly was then reviewed using 3d-dna (<u>Dudchenko et al., 2017</u>), according to manual adjustment. Using the modified 3d-dna and Juicebox workflow, 12 chromosomes were anchored. We obtained an initial 390.72 Mb Acuce genome assembly from 46.84 Gb PacBio single-molecule sequencing data with contig N50=5.67 Mb. After polishing the assembly, we anchored 385.15 Mb (98.43%) sequences into 12 pseudo-chromosomes using 60.84 Gb Hi-C data.

### RNA sequencing

An Iso-Seq library for leaf tissue was prepared according to the Isoform Sequencing protocol (Iso-Seq) using the Clontech SMARTer PCR cDNA Synthesis Kit and the BluePippin Size Selection System protocol as described by Pacific Biosciences (PN 100-092-800-03). The library was then sequenced using the PacBio Sequel System. The sequenced data were managed using the SMRT Analysis software suite (PacBio, Version: release_6.0.0.47841), resulting in a set of full-length transcripts.

### Repeat and gene annotation

Tandem repeats across the genome were identified using Tandem Repeat Finder (TRF, Version: 4.09) (<u>Benson, 1999</u>). Transposable elements (TEs) were detected using both homology-based methods, including RepeatMasker (Version: open-4.0.9) (<u>Chen, 2004</u>) against Repbase (<u>Jurka et al., 2005</u>) (Repbase Release 20181026; http://www.girinst.org/repbase/index.html) and de novo approaches such as LTR FINDER (Version: 1.07) (<u>Xu and Wang, 2007</u>) and ltrharvest in GenomeTools (Version: 1.5.10) (<u>Gremme et al., 2013</u>). Results from LTR_retriever (Version: 2.8) (<u>Ou and Jiang, 2018</u>) and RepeatModeler (Version: 2.0, http://repeatmasker.org/) were integrated to enhance TEs predictions.

Gene annotation was consolidated using EVidence Modeler (EVM) (<u>Haas et al., 2008</u>), combining predictions from Augustus (Version: 2.5.5) (<u>Stanke et al., 2008</u>), Genescan (Version: 2015-10-31) (<u>Burge and Karlin, 1997</u>), GlimmerHMM (Version: 3.0.2) (<u>Majoros et al., 2004</u>), and SNAP (Version: 2006-07-28) (<u>Korf, 2004</u>) with homology-based annotations derived from protein sequences of related *Oryza* species to consolidate ab initio gene predictions and homologous method annotations into a final gene set. For ab initio gene prediction, Augustus (Version: 2.5.5) (<u>Stanke et al., 2008</u>) was run with the trained model of rice. Three additional ab initio gene prediction software were also used: Genescan (Version: 2015-10-31) (<u>Burge and Karlin, 1997</u>), GlimmerHMM (Version: 3.0.2) (<u>Majoros et al., 2004</u>), and SNAP (Version: 2006-07-28) (<u>Korf, 2004</u>). Protein sequences of the accessions Nipponbare (<u>Goff et al., 2002</u>), 93-11 (<u>Yu et al., 2002</u>), Shuhui498 (R498) (<u>Du et al., 2017</u>), *Oryza barthii (*<u>Jacquemin et al., 2013</u>), *O. glaberrima* (<u>Wang et al., 2014</u>), and *O. nivara* (<u>Jacquemin et al., 2013</u>) species were obtained and used for homology-based gene annotation. GeneWise (Version: 2.2.0) (<u>Birney and Durbin, 2000</u>) was then used to predict gene structure within each protein-coding region. Based on all the above-mentioned annotation results, a weighted and non-redundant gene set was generated by merging all of the gene models with Acuce A15 EVM (Version: r2012-06-30625). Finally, the full-length transcripts were fed into PASA pipeline (Version: 2.4.1) (<u>Haas et al., 2008</u>) together with the EVM results for gene structure refinement and alternatively spliced isoform annotation. The genes were filtered if the proportion of repeat sequences in the gene sequences was greater than 40% or if the CDS sequence length was less than 150 bp. Gene functions were assigned according to the best match alignment using blastp against KEGG databases. InterProScan functional analysis and Gene Ontology IDs were obtained using InterProScan (Version: 5.53-87.0) (<u>Zdobnov and</u> <u>Apweiler, 2001</u>).

The pathways to which the genes belonged were derived from the matching genes in KEGG. The completeness of predicted genes was assessed using BUSCO (Version: 5.4.4) (<u>Simão et</u> <u>al., 2015</u>) with embryophyte_odb10 lineage dataset (2020-09-10).

### AIGs and NLRs comparisons

The pairwise gene synteny between the R498 reference genome and Acuce line 15 was identified using MCscan (Python version, JCVI package, Version: 1.2.7) (<u>Tang et al., 2008</u>). AIGs were then extracted from <u>Wang et al. (2018)</u>. NLRs were identified using full-length Homology-based R-gene Prediction (HRP) pipeline (<u>Andolfo et al., 2022</u>). Because NLRs are known to be highly variable, SNP calling was performed with the Acuce genome as reference according to above method. The nucleotide heterozygosity of each SNP was computed using VCFtools software (Version: 0.1.13) (<u>Danecek et al., 2021</u>). The nucleotide diversity of a gene was calculated as nucleotide divergence of all SNPs in the gene region divided by gene length. Gene presence/absence variation was characterized using the SGSGeneLoss package (Version: 0.1) (<u>Golicz et al., 2015</u>) based on above sorted BAM files. For a given gene in a given accession, if less than 20% of its exon regions were covered by at least two reads (minCov = 2, lostCutoff = 0.2), this gene was treated as absent in that accession, otherwise it was considered present.

### Search for positive selection

SweeD (Version: 3.3.1) (<u>Pavlidis et al., 2013</u>) was applied to SNP matrix to detect selective sweeps based on the CLR test to detect signatures of artificial or natural selection in each rice group with grid number set as chromosome length / 1000. A 50 kb sliding window approach was applied to quantify polymorphism levels (*π*, pairwise nucleotide variation as a measure of variability) by using VCFtools software (Version: 0.1.13) (<u>Danecek et al., 2021</u>). The impact of NLRs presence on *π* was assessed using a Wilcoxon test. The fold change was calculated between the average *π* of regions containing NLRs and those without. These analyses were conducted on both populations, Acuce and XI, for the whole genome and each chromosome. The CLR threshold value was set to select the 0.5%.

### Balancing selection

Balancing selection was identified using Beta2 statistic in BetaScan (<u>Siewert and Voight,</u> <u>2020</u>) applied to SNPs matrix with a sliding window size of 1000 bp, the substitutions and ancestral alleles were called using *O. rufipogon* as outgroup. The mutation rate was calculated as 4 × Ne × µ based on the site frequency spectrum of SNPs in putatively neutrally evolving regions of the genome, where µ is the locus neutral mutation rate, Ne is the effective population size. The threshold for Beta2 score was set as top 1%.

### GWAS analysis

GWAS was performed on rice brown spot using field data. BLUP (best linear unbiased predictors) were calculated using a generalized linear mixed model implemented in the glmer function from the R package lme4 (Bates et al., 2015), with a logit link function and a binomial distribution of errors:

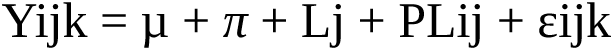

where Yijk was the severity of the rice brown spot from plot i, line j and repetition k, *π* was the random effect of the plot i, Lj was the random effect of the line j, and PLij was the interaction between the plot i and the line j. A GWAS was then performed with the BLUP values based on SNP set called with Acuce genome using Multiple Loci Mixed linear Model (MLMM) in GAPIT (<u>Lipka et al., 2012</u>). The first five PCA values (eigenvectors) were used as fixed effects in the model to correct for stratification. Considering that 27,522 SNPs were used, we defined the whole-genome significance cutoff with -log10 (P) = 5.

## Results

### Acuce samples display significant variation for pathogen susceptibility but not for agronomical traits

We first examined the distribution of phenotypic traits expressed by Acuce samples in the field, considering agricultural traits as well as disease resistance. This analysis was conducted using the subset of 89 inbred lines. We observed negative correlations between, on one hand, levels of damages caused by the two main rice pathogens (blast and smut fungi) and, on the other hand, thousand-seed weight, and yield (Figure S2), suggesting that fungal pathogens are potential drivers of plant fitness. This is consistent with current knowledge that pathogens are major, worldwide determinants of plant yield (<u>Savery et al., 2019</u>). As a whole, major agronomic traits such as plant height, tiller number, and thousand-seed weight exhibited low variability in Acuce inbred lines (Figure 1A, B). With respect to these agronomic traits, levels of disease resistance to three major fungal pathogens monitored were more variable. This high variability was also observed under controlled conditions for blast (Table S1), supporting the hypothesis that the observed variability of resistance in the field is due to genetic variation and not to heterogeneous levels of pathogen pressure. Thus, although major traits easily tractable by farmers were quite homogeneous in Acuce, disease resistance traits appeared to be highly variable.

**Figure 1.**
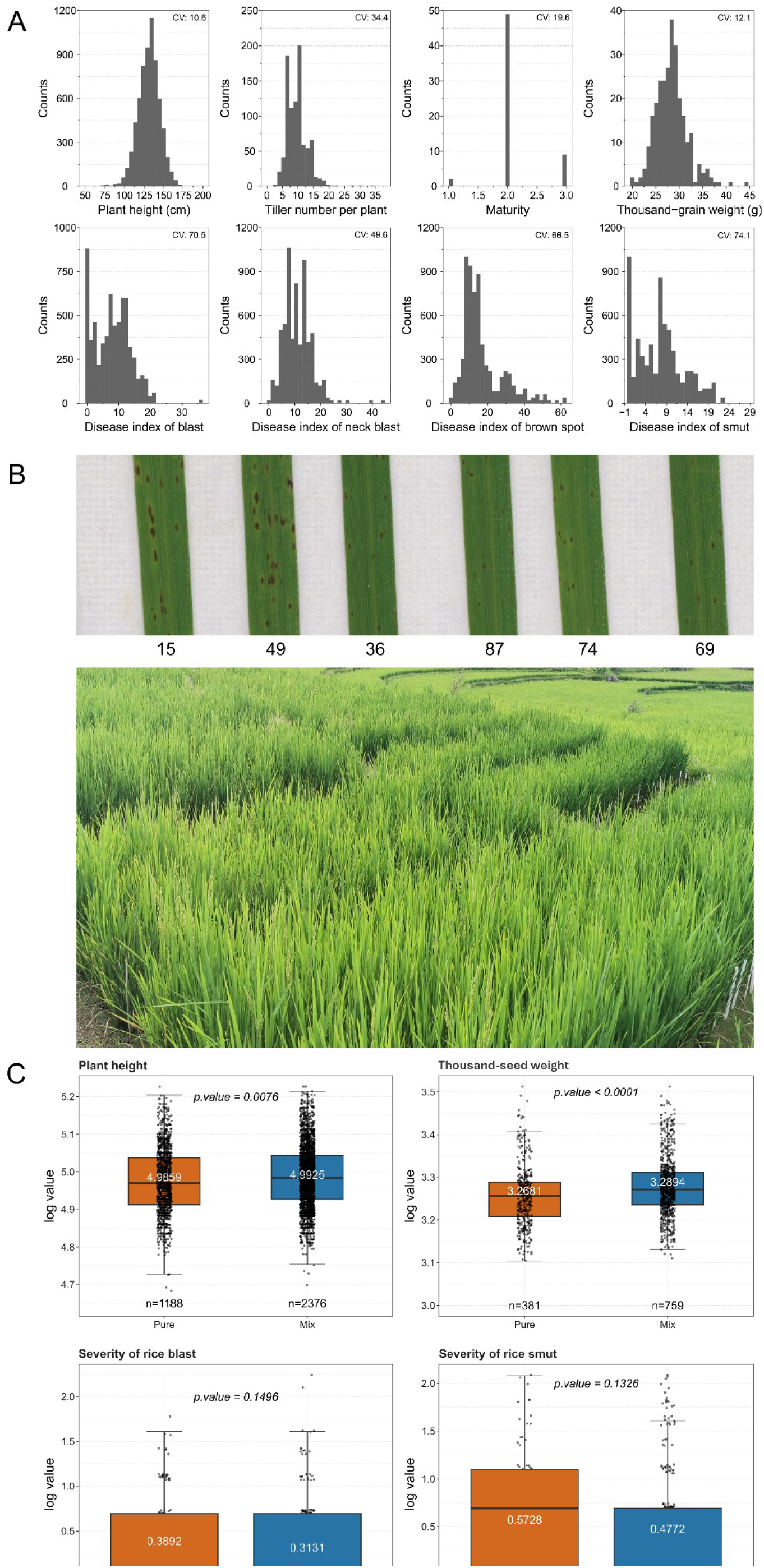
Phenotypic diversity in the Acuce landrace and emerging properties in binary mixtures. Field phenotypes of 89 inbred Acuce lines grown in pure stands (A and B) or in binary mixtures (C). (A) Distribution of growth phenotypes are weakly variable (coefficient of variation values are indicated on each panel) compared to disease phenotypes. (B) Examples of Acuce lines grown in the field displaying some phenotypical differences (forefront) and paddy fields in the background showing the homogeneity of the landraces grown by farmers. The upper panel shows representative symptoms of brown spot under laboratory conditions with the isolate CH9015 used for GWAS analysis (Figure S6). (C) Six binary mixtures of selected Acuce inbred lines were phenotyped in the field for three years. The average values of severity in pure stand and mixtures are shown. Significant differences were evaluated using *t*-test for plant height or thousand-seed weight and Wilcox test for incidence of disease.

### Mixing different genotypes results in superior performances in the field

Under field conditions, we tested the hypothesis that emerging phenotypes in mixtures of Acuce inbred lines could favour the maintenance of genetic diversity by increasing the fitness of the population. Artificially reconstituting diversity by combining two-by-two Acuce inbred lines weakly (0.1 to 0.7%) but significantly increased plant height and thousand-seed weight (Figure 1C), with some mixtures being more effective than others (Figure S3B). Mixtures tended to reduce disease levels: the incidence of smut (number of plants showing symptoms) was reduced by 23% in mixtures compared to pure stands (Figure S3D), and the average severity (defined as the level of symptoms) was also reduced by 17% (Figure 1C). Due to discretization of source data and smaller dataset size, this result not significant overall. However, it was significant in certain pairs of Acuce lines, with a severity reduced by up to 62% in some cases (Figure S3C). Although disease suppression, in particular incidence levels, is expected in cultivar mixtures (<u>Mundt, 2002</u>), it usually requires a minimum size of the plots to be observed, which was not the case in our experimental design consisting of small plots. Rather, we propose that the mere mixing of Acuce lines generates emergent properties, at the plant level and measured at the severity level, on traits that are key components for yield. This is consistent with observations made under controlled conditions where disease severity to pathogens was modulated by plant-plant interactions in Acuce mixtures (<u>Pélissier et al.,</u> <u>2023</u>).

### Population structure at the genome level

The genomes of 227 Acuce (including 89 Acuce inbred lines) and 30 HY3 samples were compared to representative genomes of 192 YYT and 50 worldwide XI accessions (Table S1; Figure S1E). We identified a total of 20,731,390 single nucleotide polymorphisms (SNPs) that were considered for analysis. To reduce the computational burden, only 27,522 representative sites were used for structure analysis. The optimization criterion suggested that K=9 was the most appropriate for biological interpretation (Figure 2A, Figure S4) allowing the identification of three sets of Acuce samples (with each including inbred lines) belonging to different clusters (hereafter called “families”; Table S1; Figure S4C). Phylogenetic analysis grouped accessions according to their cluster assignation (Figure 2B). Acuce accessions were dominant in clusters A and I while many others belonged to cluster E, mixed with many non-Acuce YYT1 accessions. The rest of Acuce accessions, all of them originating from a single village (Tur Guo Zhai), belonged to cluster F, along with YYT1 accessions and one HY3 sample. We ignored this group of accessions because the single-village origin was indicative of a potential labelling error, and we defined the three families of Acuce accessions after the assignation to the A, I, and E clusters. HY3 formed a unique, different group, in agreement with its known pedigree (Figure S1C). In the phylogenetic tree, the three Acuce families and the YYT4 cluster were imbedded into the YYT1 cluster, and Acuce accessions were included into both YYT1 and YYT4-dominated clades, in agreement with <u>Huang et al. (2025)</u> who found Acuce accessions in both these clusters. Using structure analysis and phylogenetic trees, we discarded samples labelled as Acuce that clearly clustered with non-Acuce landraces as they may result from mislabelling or seed contamination which is common in the YYT (<u>Huang et al., 2025</u>). Similarly, we excluded samples labelled as HY3 that were not associated with the relevant cluster. As a result, we defined conservative groups containing 108 “true” Acuce (excluding inbred), 87 “true” inbred Acuce, and 23 “true” HY3 (Figure S4D). The following analyses were based on these three groups and two sets of reference samples: worldwide XI samples and samples from other YYT landraces (denoted “YYT”).

**Figure 2.**
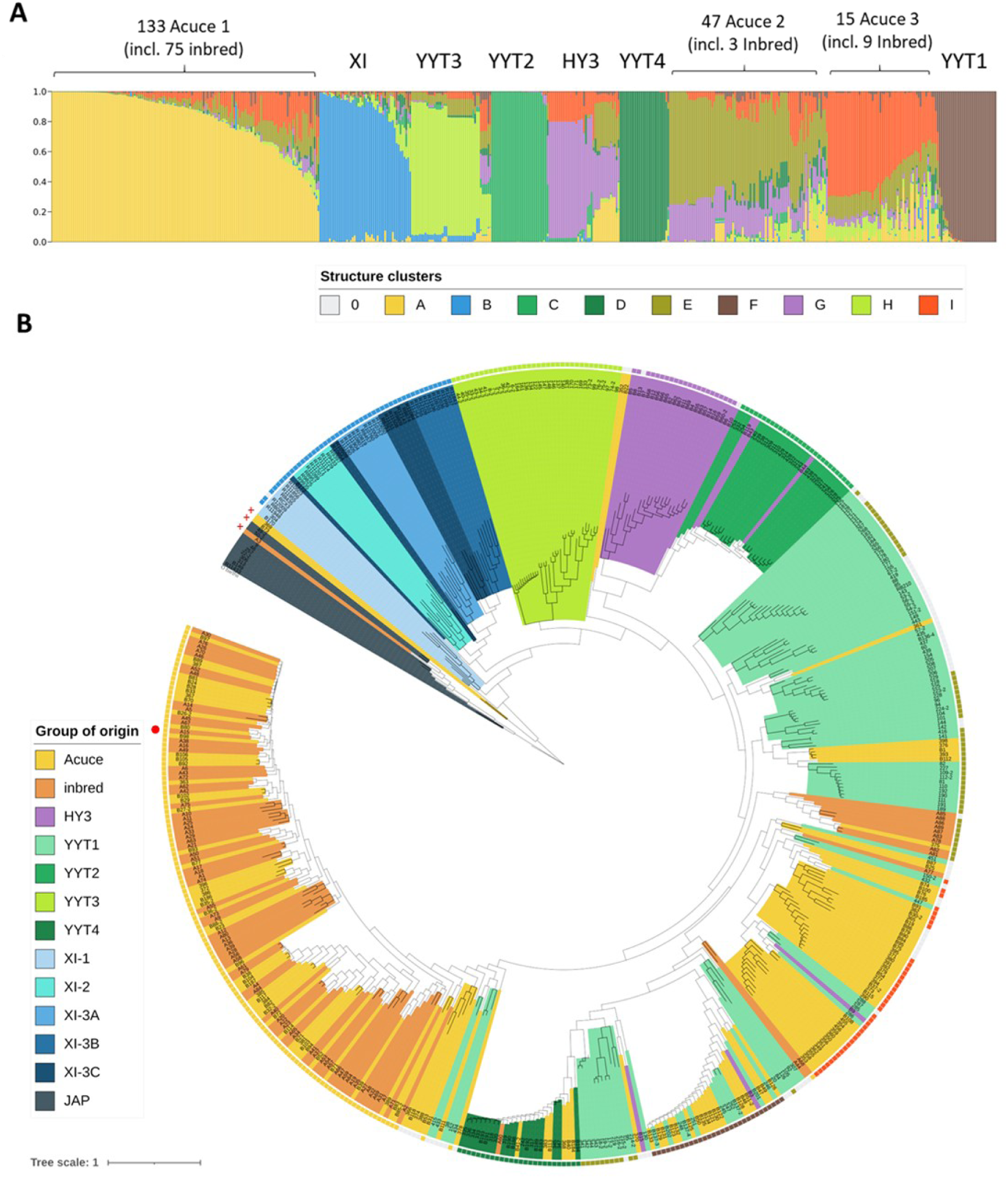
Structure of genetic diversity in Acuce landrace and its relatives. (A) The top panel represents the assignment probability to the clusters of the structure analysis of 494 indica samples (K=9). Cluster assignment was used to define true Acuce and delineate Acuce families 1 to 3. When no assignment was possible, the position in the tree was considered. (B) The distance-based tree contains 510 samples (including 10 confirmed and 5 potential non-indica samples, and the outgroup *Oryza barthii*). Branch lengths are square-root scaled for better readability. Clades are colored according to the group of origin of samples, and outer markings indicate assigned cluster in the structure analysis. The worldwide XI (XI-1 to XI-3C) and GJ (JAP) sub-species as defined in <u>Wang et al. (2018)</u> are indicated as well as four sub-groups of YYT landraces identified in separate studies (<u>Ali et al., 2023</u>; <u>Huang et al.</u>). Red “+” signs represent probable seed contaminations of Acuce fields by genotypes detected by extreme genetic divergence from the bulk of Acuce accessions. The red dot indicates the Acuce line A15 used for Pacbio sequencing. A15

### Levels of nucleotide diversity

The comparison of diversity levels between groups showed that diversity within Acuce was as high as 38% of the levels found in the entire XI subspecies (Table S2), consistently with previous studies that showed that landraces harbour high levels of diversity in YYT (<u>Gao et</u> <u>al., 2012</u>). On average, nucleotide diversity values were 28% higher in Acuce than in inbred and 71% higher than in the recently introduced HY3 population. Quite remarkably, the range of differentiation values (estimated with the Jost’s D) between Acuce families overlapped those between worldwide XI clusters (Figure S1B and Table S5). There was no strong difference in nucleotide diversity among Acuce families when considering villages or clusters of villages (Table S2). Considering villages with sufficient sample sizes, the differentiation varied from 1.1% to 6.3%, much below differentiation between Acuce families (Table S2 reported in Figure S1B). This small difference between villages is consistent with the strong social habit of exchanging seeds in YYT (<u>Dedeurwaerdere and Hannachi, 2019</u>).

### Genome-wide signatures of selection

We performed scans for signatures of selection (either natural or artificial). Pervasive selective sweeps were detected along the genome using SweeD for both Acuce (including inbred), and HY3 groups, targeting genes encoding proteins belonging to a variety of functional categories (Tables S3 and S4). The CLR cut-off values selecting the top 0.5% of regions were as follows: 1795.98 for Acuce family 1, 3263.83 for Acuce family 2, 1725.57 for inbred family 1, 2708.91 for inbred, 1639.96 for Acuce and 2540.68 for HY3. Interestingly, there was little overlap between the functional categories and genes detected in the two populations. This is expected if the two varieties underwent independent breeding histories (Figure S5).

We identified genes targeted by balancing selection using BetaScan. We found a significant association between genes targeted by balancing selection and gene ontology only in the case of Acuce. All identified ontologies were associated with Nucleotide-binding domain and Leucine-rich Repeat genes (NLRs; Table S6) which are major determinants of pathogen resistance (<u>Ngou et al., 2022</u>). The mode of evolution of NLRs, involving in particular extensive presence/absence variation might blur the signal. Therefore, we performed a specific analysis of NLRs diversity using an *ad hoc* genome reference from one of the Acuce samples using PacBio sequencing (Table S7 and Figure S7). We identified 452 NLRs in this *de novo* Acuce genome (Table S8), a rather small number compared to other known rice genomes (e.g. 712 NLRs were identified in the R498 genome) but consistent with the analysis of <u>Gladieux et al. (2024)</u> who identified 460 NLRs in Acuce. Thirteen NLRs were identified under balancing selection in Acuce, 4 times more than expected by chance (Table S6). None of these NLRs corresponded to known resistance genes, suggesting that Acuce landrace achieves durable resistance with a yet unknown set of rapidly evolving resistance genes. Besides, we performed a GWAS analysis based on field phenotyping data with respect to resistance to several common diseases affecting rice. We identified two SNPs on chromosome 2 nearby a gene coding for a Leucine-rich repeat protein associated with brown spot resistance (Figure S6). This result still needs to be confirmed with further study, but this finding demonstrates the potential of Acuce as a resource for the identification of determinants of disease resistance. Quite interestingly, the recently introduced HY3 modern population also showed some NLRs under balancing selection. This suggests that this pattern of selection is connected to cultural practices applied in the YYT, from where HY3 samples have been collected.

### Generation and annotation of a Acuce reference genome

A *de novo* genome assembly for Acuce accession A15 was generated in order to provide a relevant reference for surveying presence-absence variation of NLR genes in Acuce samples (Table S12). The annotation procedure achieved 98.2% completeness of the BUSCO core embryophyta genes (Table S12c). We predicted ∼215.41 Mb (55.05%) repetitive elements in the Acuce genome (Table S12d), among which 212.86 Mb repeats were transposable elements (TEs; Table S12e). We predicted 40,011 protein-coding genes (41,619 transcripts) following a combination of homology and ab initio methods (Table S12f), which captured 96.8% complete BUSCOs in embryophyte_obd10 (Table S12g). 67% (26,803) genes were assigned to at least one function in InterPro, GO or KEGG (Table S12h).

### Genomic diversity provides Acuce with diverse immunity factors

We compared the diversity of NLRs to other agronomically important genes (AIGs) thanks to the *de novo* A15 genome. AIGs involved in leaf architecture, tillering and flowering were identified by database homology in the Acuce genome (Table S9). Fifty-seven of them were unambiguously found in Acuce according to the gene synteny and blastp check (Table S9). Overall, NLRs were much more diverse than AIGs in Acuce, inbred Acuce and HY3 (Figure 3A; Table S8). In addition to single polymorphic sites, we looked for presence/absence variations: Acuce had an average of 446 NLRs with a variance of 13.7 (449 for inbred, variance 8) as compared with 435 with a variance of 11.3 for HY3 (Figure 3B; Table S10). There was therefore a slightly elevated number of NLRs in Acuce when compared with HY3. Typical feature of diversity landscape for the Acuce landrace and XI genomes is shown in Figure 3C and Figure 3D, showing large-scale variation in both levels of diversity and presence of NLRs (Figure S8, Table S13). A significant association of diversity with regions containing NLRs was found for both populations (Figure 3E), with a stronger enrichment in Acuce (fold change=1.51) compared to XI genomes (fold change=1.37), for some but not all chromosomes (Figure 3F).

**Figure 3.**
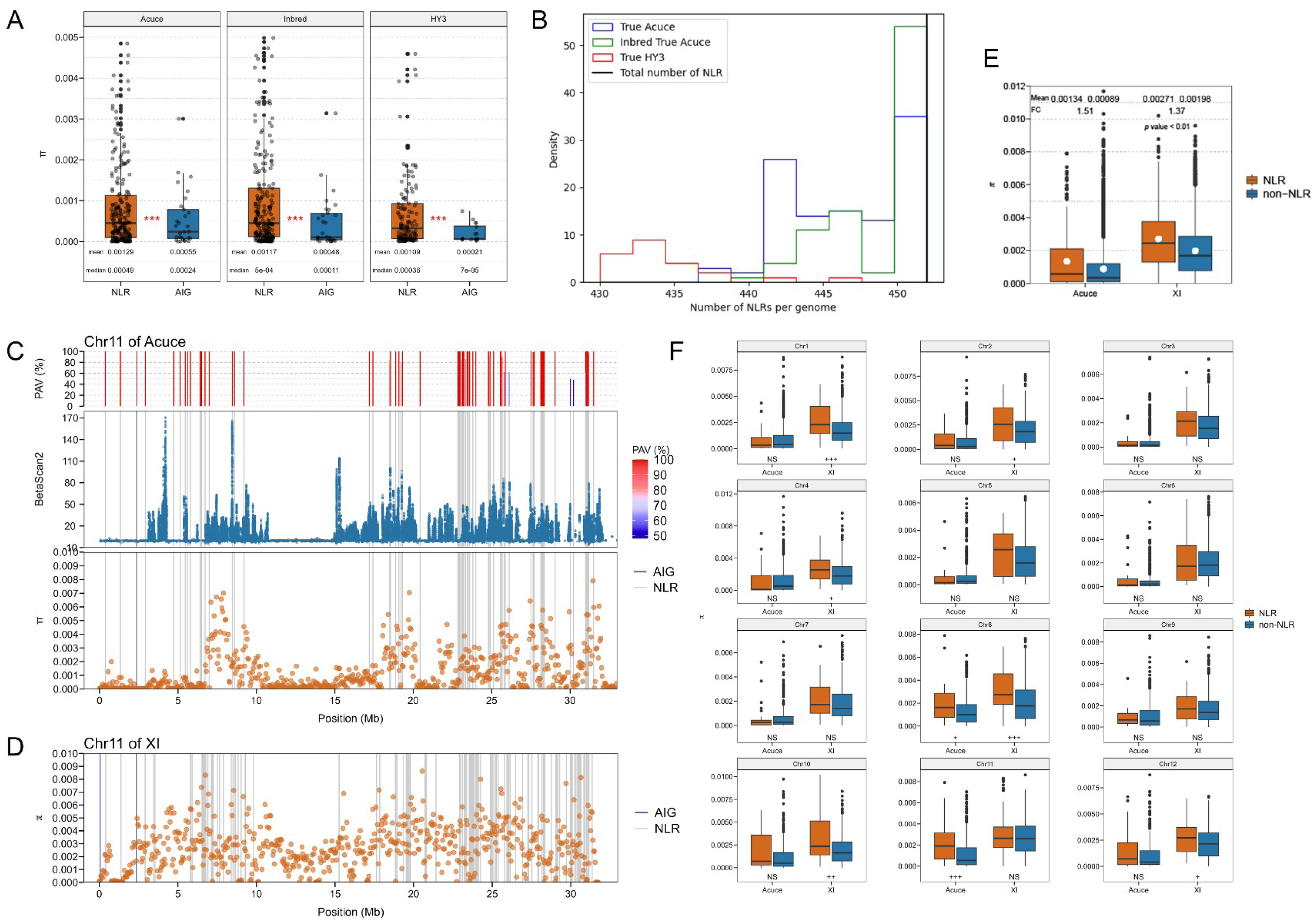
Genomic diversity landscape of NLR and agronomically important genes in Acuce landrace. The average diversity levels (π) of NLRs and AIGs (A) and the distribution of NLR genes (B) were analyzed in 108 true Acuce, 87 true inbred Acuce and 23 true HY3 genomes. The red plus signs (+++) signify statistical significance at p < 0.001 based on the Wilcoxon Rank Sum and Signed Rank Tests. (C) The average diversity (π) and balancing selection (betascan2) levels (50 kb window) are shown along the chromosomes 11 of the Acuce line 15 genome, as well as the position of NLR and AIGs. Their average presence/absence variation (PAV) in the 108 true Acuce genomes is also displayed. (D) The average diversity (π) levels (50 kb window) are shown along the chromosomes 11 of the R498 genome. (E) For both Acuce and XI, average diversity levels (π) of NLR and non-NLR bins were calculated within 50 kb windows. The term “mean” represents the average value, “fold change” indicates the multiplicative change in the mean values of NLR/non-NLR, and the p-value is computed using the Wilcoxon test. (F) On each chromosome, the average diversity (π) levels of NLR and non-NLR bins were computed within 50 kb windows for both Acuce and XI populations. Statistical analysis was performed using the Wilcoxon test, where “NS” denotes not significant, “+” indicates a p-value < 0.05, “++” signifies a p-value < 0.01, and “+ ++” represents a p-value < 0.001.

## Discussion

Landraces are considered to be untapped reservoirs for future breeding efforts (<u>McCouch et</u> <u>al., 2020</u>). They have long been known to be more successful than modern cultivars under stressful conditions (<u>Dwivedi et al., 2016</u>; <u>Newton et al., 2010</u>). Our in-depth genomic analysis of Acuce showed that, despite visible homogeneity for agricultural traits likely under direct stabilizing selection by farmers, very high levels of genomic diversity were maintained within this landrace. This diversity is specifically inflated in immunity receptors, resulting in high phenotypic variation in disease resistance traits, consistenly with patterns observed for other landraces cultivated in the YYT (<u>Gladieux et al., 2024</u>). The analysis of HY3 indicates that modern breeding can result in substantial levels of diversity, albeit at a lower level than those observed in the traditional Acuce landrace. Through signatures of balancing selection and the results of GWAS analysis, we provide several lines of evidence hinting that this diversity allows Acuce plants to respond to selection imposed by pathogens, similarly to other YYT landraces (<u>Huang et al., 2025</u>).

The high levels of diversity observed within Acuce can be traced back to the origins of the landrace and may have been maintained continuously by agricultural practices or, alternatively, might be fuelled by recurrent or permanent gene flow, as suggested by the sharing of ancestry groups with other landraces. Deciphering the origins of Acuce diversity requires further study. Our results clearly point to strong levels of artificial stabilizing selection over agricultural traits, both at the genomic and phenotypic levels. In contrast, selection on disease resistance did not exhaust polymorphism of resistance genes. Crop protection against pathogens may be under direct selection by farmers in Acuce, but heterogeneous disease incidence (in both time and space) might prevent complete fixation of single alleles at resistance loci, ultimately allowing for the coexistence of different alleles at immunity receptor genes.

Our results show that, in the field, several traits related to yield, which in some respect may be considered as fitness, are improved by simply mixing two genomes. Diversity at genotypic level has been shown to reduce disease risks and prevalence in agro-systems (<u>Mundt, 2002</u>). For instance, mixing two varieties has been proven to confer resistance to rice blast fungus (<u>Zhu et al., 2000</u>). The combination of two or more resistance genes in separate varieties has been classically invoked to explain this property and the sustainability of such agro-systems (<u>Borg et al., 2018</u>; <u>Garrett et al., 2009</u>). To some extent, disease control was also reduced in some pairwise mixtures tested here. This observation reinforces the hypothesis that direct plant-plant interactions can modulate disease resistance of individual plants (<u>Pélissier et al.,</u> <u>2023</u>; <u>Pélissier et al., 2021</u>) and thus add another possible level of disease regulation in cultivar mixtures besides known processes (<u>Borg et al., 2018</u>). Our experimental design only considered pairwise mixtures, still much simpler than the complex assemblages observed in field condition. In particular, the fitness benefit is likely to depend on the matching between, on the one hand, immunity receptor alleles expressed by genotypes composing the mixture and, on the other hand, the actual presence of corresponding pathogen genotypes. The recent release of the HY3 modern composite population, which is depleted of a dozen immune receptors, is an opportunity to challenge the stability of disease control achieved by modern populations in the field. In conclusion, the Acuce landrace exhibits extensive genomic diversity, especially in immune receptors, offering potential advantages in disease resistance and crop protection.

## Acknowledgements

We thank Elisabeth Fournier and Henry Adreit for their support in collecting plants in the field. This study was supported by National Key R&D Program of China (2023YFE0107500, 2023YED1400800), National Natural Science Foundation of China (31801792, 31960554), Yunnan Fundamental Research Projects (202401AS070002) and Yunnan Xingdian Talent Support Program. It was conducted under the MoBiDiv (https://www6.inrae.fr/mobidiv, funded by the ANR program PPR-CPA ANR-20-PCPA-0006), the SuMCrop INRAE Metaprogram, the ANR MUSE (ANR-16-IDEX-0006 AMUSER) and the INRAE-YAU LIA PLANTOMIX projects.

## Author contributions

HH and JBM conceived and designed the study. CL and YZ collected the genetic material. HH, XL and LM processed the samples and performed the DNA extraction. XL, LM, RP, JL and JL phenotyped the inbred lines in the field. YZ and XH contributed to the historical understanding of Yuanyang terraces. YD generated the original sequencing data. SDM, SD, XL, RP, HH and JBM analyzed the data and prepared the figures. LM helped for GWAS analysis and statistical analysis. SDM, SD, XL, YD, HH and JBM wrote the manuscript draft, the co-authors reviewed and edited the manuscript. All authors read and approved the final manuscript.

## Data availability

SNP tables of all individuals sequenced in this study after mapping to the two reference genomes are available through an open web server: - http://teabase.ynau.edu.cn/data/tea/pangeome/rice/Table_S12.vcf.gz, - http://teabase.ynau.edu.cn/data/tea/pangeome/rice/Table_S14.vcf.gz.

## Supporting information

**Fig. S1 Origins of Acuce and HY3 genomes.**

**Fig. S2 Field phenotypes of Acuce inbred lines.**

**Fig. S3 Phenotypes in binary Acuce mixtures.**

**Fig. S4 Structure analysis of Acuce and HY3 genomes.**

**Fig. S5 Separate breeding histories between Acuce and HY3.**

**Fig. S6 GWAS analysis of brown spot resistance in the Acuce landrace.**

**Fig. S7 Heatmap of Hi-C interactions for Acuce pseudo-chromosomes.**

**Fig. S8 Diversity feature of Acuce and XI genomes.**

**Table S1 Origin and associated metadata for all rice lines used in this study.**

**Table S2 Diversity statistics per population of Acuce and other local and indica genomes.**

**Table S3 Genes under purifying selection in true Acuce, Indred and HY3 sub-groups.**

**Table S4 Gene ontology enrichment for genes under purifying selection.**

**Table S5 Pairwise comparisons of Auce and other local and indica genomes.**

**Table S6 Genes under balancing selection in true Acuce, Indred and HY3 sub-groups.**

**Table S7 Genome annotation of the Acuce line 15, including NLR genes.**

**Table S8 Nucleotide diversity at AGI and NLR genes.**

**Table S9 Agronomically important genes (AIG) in Acuce genome.**

**Table S10 Presence/Absence polymorphism for AIG and NLR genes in Acuce and HY3.**

**Table S11 Genome statistics.**

**Table S12 Statistics of the Acuce genome assembly. Table S13 Table for average diversity.**

