## Supplementary Material for "Population genomics of the rice landrace Acuce reveals exceptional dynamics of immune receptors"

**Fig. S1.** Origins of Acuce and HY3 genomes.

**Fig. S2.** Field phenotypes of Acuce inbred lines.

**Fig. S3.** Phenotypes in binary Acuce mixtures.

**Fig. S4.** Structure analysis of Acuce and HY3 genomes.

**Fig. S5.** Separate breeding histories between Acuce and HY3.

**Fig. S6.** GWAS analysis of brown spot resistance in the Acuce landrace.

**Fig. S7.** Heatmap of Hi-C interactions for Acuce pseudo-chromosomes.

**Fig. S8.** Diversity feature of Acuce and XI genomes.

**Table S1.** Origin and associated metadata for all rice lines used in this study.

**Table S2.** Diversity statistics per population of Acuce and other local and indica genomes.

**Table S3.** Genes under purifying selection in true Acuce, Indred and HY3 sub-groups.

**Table S4.** Gene ontology enrichment for genes under purifying selection.

**Table S5.** Pairwise comparisons of Auce and other local and indica genomes.

**Table S6.** Genes under balancing selection in true Acuce, Indred and HY3 sub-groups.

**Table S7.** Genome annotation of the Acuce line 15, including NLR genes.

**Table S8.** Nucleotide diversity at AGI and NLR genes.

**Table S9.** Agronomically important genes (AIG) in Acuce genome.

**Table S10.** Presence/Absence polymorphism for AIG and NLR genes in Acuce and HY3.

**Table S11.** Genome statistics.

**Table S12.** Statistics of the Acuce genome assembly.

**Table S13.** Table for average diversity.

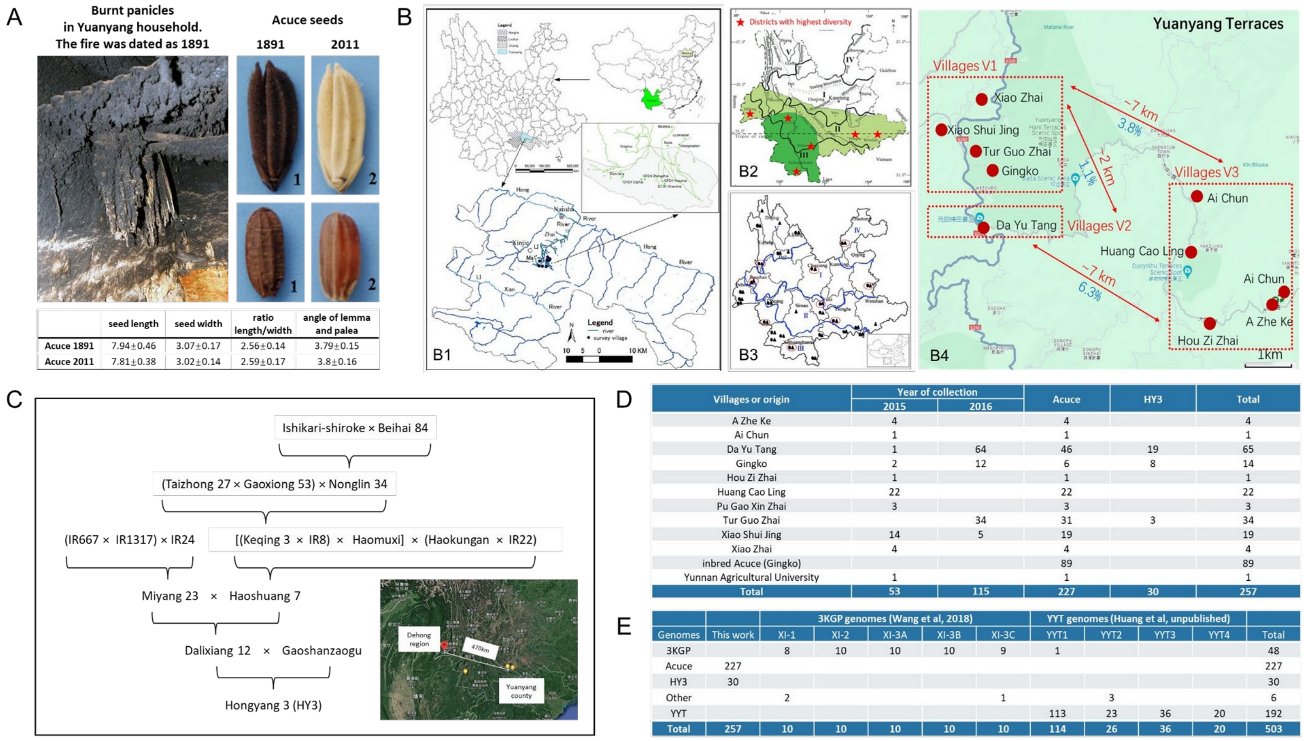


| | seed length | seed width | ratio length/width | angle of lemma and palea |
|---|---|---|---|---|
| Acuce 1891 | 7.94±0.46 | 3.07±0.17 | 2.56±0.14 | 3.79±0.15 |
| Acuce 2011 | 7.81±0.38 | 3.02±0.14 | 2.59±0.17 | 3.8±0.16 |

| Villages or origin | Year of collection | | Acuce | HY3 | Total |
|---|---|---|---|---|---|
| | 2015 | 2016 | | | |
| A Zhe Ke | 4 | | 4 | | 4 |
| Ai Chun | 1 | | 1 | | 1 |
| Da Yu Tang | 1 | 64 | 46 | 19 | 65 |
| Gingko | 2 | 12 | 6 | 8 | 14 |
| Hou Zi Zhai | 1 | | 1 | | 1 |
| Huang Cao Ling | 22 | | 22 | | 22 |
| Pu Gao Xin Zhai | 3 | | 3 | | 3 |
| Tur Guo Zhai | | 34 | 31 | 3 | 34 |
| Xiao Shui Jing | 14 | 5 | 19 | | 19 |
| Xiao Zhai | 4 | | 4 | | 4 |
| inbred Acuce (Gingko) | | | 89 | | 89 |
| Yunnan Agricultural University | 1 | | 1 | | 1 |
| Total | 53 | 115 | 227 | 30 | 257 |

| | | 3KGP genomes (Wang et al, 2018) | | | | | YYT genomes (Huang et al, unpublished) | | | | |
|---|---|---|---|---|---|---|---|---|---|---|---|
| Genomes | This work | XI-1 | XI-2 | XI-3A | XI-3B | XI-3C | YYT1 | YYT2 | YYT3 | YYT4 | Total |
| 3KGP | | 8 | 10 | 10 | 10 | 9 | 1 | | | | 48 |
| Acuce | 227 | | | | | | | | | | 227 |
| HY3 | 30 | | | | | | | | | | 30 |
| Other | | 2 | | | | 1 | | 3 | | | 6 |
| YYT | | | | | | | 113 | 23 | 36 | 20 | 192 |
| Total | 257 | 10 | 10 | 10 | 10 | 10 | 114 | 26 | 36 | 20 | 503 |

**Fig. S1. Origins of Acuce and HY3 genomes.**

All lines studied here were isolated from individual panicles. **A.** The seeds found in the burnt household (dated 1851) are similar to current Acuce seeds according to morphological characters. Unfortunately, DNA could not be extracted from these seeds. **B.** The area where seeds were collected are compared to the areas reported in previous studies (**B1**: Jiao et al., 2012; **B2**: Zhang et al, 2007; and **B3**: Cui et al, 2019). B4. Areas of collection of Acuce and HY3 seeds. **C.** Pedigree of the HY3 variety. It incorporates genomes from Yunnan Dehong region (Haomuxi and Haokungan), from YYT (Gaoshanzaogu) and from Taiwain (Keqing 3) or other modern varieties (IR accessions). **D.** Summary table of origins (village and year) of the Acuce and HY3 lines. E. Summary table of the origin of all genomes used in this study.

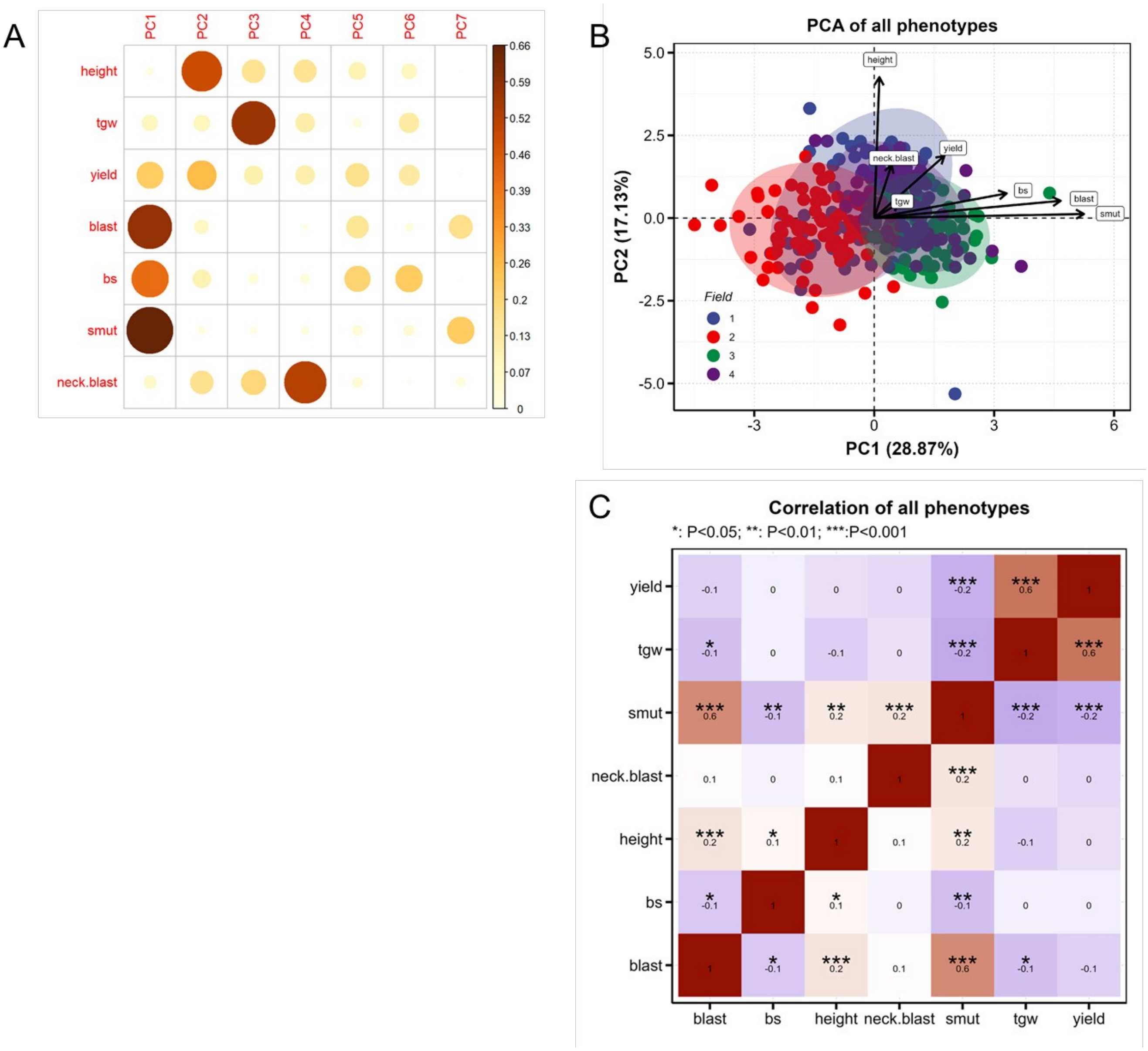
A
PC1
PC2
PC3
PC4
PC5
PC6
PC7
height
tgw
yield
blast
bs
smut
neck.blast
0.66
0.59
0.52
0.46
0.39
0.33
0.26
0.2
0.13
0.07
0
B
PCA of all phenotypes
PC2 (17.13%)
PC1 (28.87%)
5.0
2.5
0.0
-2.5
-5.0
-3
0
3
6
height
neck.blast
yield
tgw
bs
blast
smut
Field
1
2
3
4
C
Correlation of all phenotypes
*: P<0.05; **: P<0.01; ***:P<0.001
yield
tgw
smut
neck.blast
height
bs
blast
blast
bs
height
neck.blast
smut
tgw
yield

**Fig. S2. Field phenotypes of Acuce inbred lines.**

The data represent the normalized values of phenotypes collected in 2022 in four widely separated fields in the Gingko village. Foliar disease phenotypes monitored were rice blast and brown spot (bs). Panicle diseases were smut and neck blast. Growth-related phenotypes were plant height, thousand grain weight (tsg) and yield. **A.** $Cos^2$ values of all phenotypes in the first 7 axis of Principal Component analysis. PC1 and PC2 represent 45% of variability. Tgw and neck blast are most explained by PC3 and PC4 respectively. **B.** All scored phenotypes displayed on the first two axis of the Principal Component Analysis. **C.** Correlation matrix of field phenotypes. The value in each bow represents the correlation (r; Pearson) value and stars the significance of the value as evaluated using the function corAndPvalue in R package WGCNA (*P<0.05; ** P<0.01; *** P<0.01).

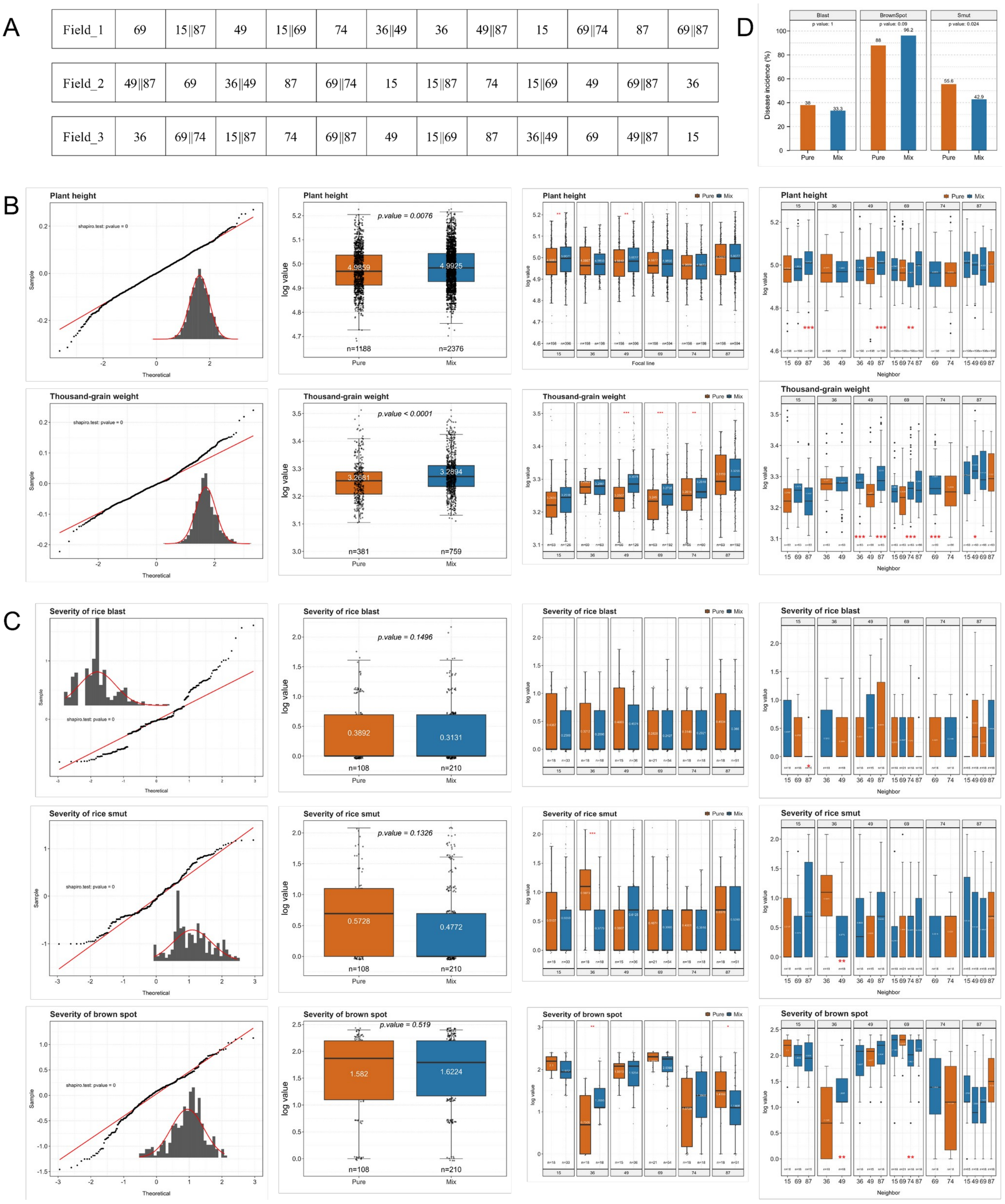
A
Field_1 | 69 | 15||87 | 49 | 15||69 | 74 | 36||49 | 36 | 49||87 | 15 | 69||74 | 87 | 69||87
Field_2 | 49||87 | 69 | 36||49 | 87 | 69||74 | 15 | 15||87 | 74 | 15||69 | 49 | 69||87 | 36
Field_3 | 36 | 69||74 | 15||87 | 74 | 69||87 | 49 | 15||69 | 87 | 36||49 | 69 | 49||87 | 15
D
Blast
p value: 1
BrownSpot
p value: 0.09
Smut
p value: 0.024
Disease incidence (%)
38
33.3
88
96.2
55.6
42.9
Pure
Mix
B
Plant height
shapiro.test: pvalue = 0
Sample
Theoretical
p.value = 0.0076
log value
4.9859
4.9925
n=1188
n=2376
Pure
Mix
Focal line
Neighbor
Thousand-grain weight
p.value < 0.0001
3.2681
3.2894
n=381
n=759
C
Severity of rice blast
p.value = 0.1496
0.3892
0.3131
n=108
n=210
Severity of rice smut
p.value = 0.1326
0.5728
0.4772
Severity of brown spot
p.value = 0.519
1.582
1.6224

**Fig. S3. Phenotypes in binary Acuce mixtures.**

Acuce lines were chosen based on their position on the phylogenetic tree in Figure 2 as representative of the Acuce largest sub-groups 1 and 3. **A.** Acuce binary mixtures were grown in the field and phenotyped for three successive years (2019, 2020 and 2021), together with the corresponding pure stands. For each phenotype, the distribution of the data is shown, the average values between pure stand and mixtures, the value of one given focal plant with all possible mixtures and the detailed values of each focal plants for all possible neighbors. **B.** Growth phenotypes. **C.** Disease phenotypes using all values, including healthy plants (noted 0). For instance smut disease severity in Acuce line 36 in the presence of Acuce line 49 is reduced by 62%. **D.** The incidence of three diseases under the Pure and Mix conditions represents the percentage of sick plants. The numbers above the bars represent the incidence, and the numbers between the two bars represent the Fisher's exact test p-values after Bonferroni correction.

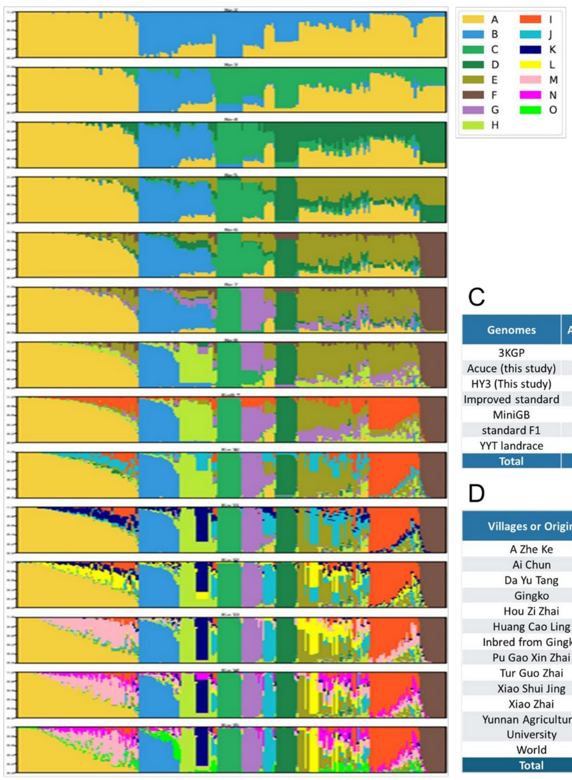


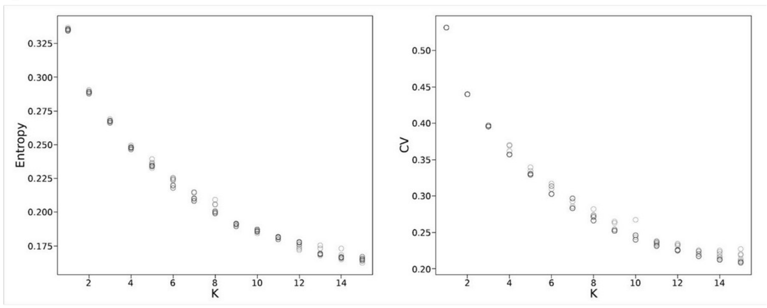


C

| Genomes | Acuce 1 | Acuce 2 | Acuce 3 | HY3 | improved | Inbred Acuce 1 | Inbred Acuce 2 | Inbred Acuce 3 | not Acuce | NOT HY3 | XI | YYT1 | YYT2 | YYT3 | YYT4 | Total |
|---|---|---|---|---|---|---|---|---|---|---|---|---|---|---|---|---|
| 3KGP | | | | | | | | | | | 48 | | | | | 48 |
| Acuce (this study) | 58 | 44 | 6 | | | 75 | 3 | 9 | 32 | | | | | | | 227 |
| HY3 (This study) | | | | 23 | | | | | | 7 | | | | | | 30 |
| Improved standard | | | | | 3 | | | | | | | | | | | 3 |
| MiniGB | | | | | | | | | | | 1 | | | | | 1 |
| standard F1 | | | | | 2 | | | | | | | | | | | 2 |
| YYT landrace | | | | | | | | | | | | 113 | 23 | 36 | 20 | 192 |
| Total | 58 | 44 | 6 | 23 | 5 | 75 | 3 | 9 | 32 | 7 | 49 | 113 | 23 | 36 | 20 | 503 |

D

| Villages or Origin | improved | Inbred True Acuce | not Acuce | NOT HY3 | True Acuce | True HY3 | XI | YYT1 | YYT2 | YYT3 | YYT4 | Total |
|---|---|---|---|---|---|---|---|---|---|---|---|---|
| A Zhe Ke | | | | | 4 | | | | | | | 4 |
| Ai Chun | | | | | 1 | | | | | | | 1 |
| Da Yu Tang | | | 2 | 3 | 44 | 16 | | | | | | 65 |
| Gingko | | | 1 | 1 | 5 | 7 | | | | | | 14 |
| Hou Zi Zhai | | | | | 1 | | | | | | | 1 |
| Huang Cao Ling | | | 1 | | 21 | | | | | | | 22 |
| Inbred from Gingko | | 87 | 2 | | | | | | | | | 89 |
| Pu Gao Xin Zhai | | | 1 | | 2 | | | | | | | 3 |
| Tur Guo Zhai | | | 21 | 3 | 10 | | | | | | | 34 |
| Xiao Shui Jing | | | 3 | | 16 | | | | | | | 19 |
| Xiao Zhai | | | 1 | | 3 | | | | | | | 4 |
| Yunnan Agricultural University | 4 | | | | 1 | | | 113 | 23 | 36 | 20 | 197 |
| World | 1 | | | | | | 49 | | | | | 50 |
| Total | 5 | 87 | 32 | 7 | 108 | 23 | 49 | 113 | 23 | 36 | 20 | 503 |

**Fig. S4. Structure analysis of 494 samples based on Admixture and sNMF.**

**A.** Assignation coefficients for each sample, averaged over the two methods and over 10 replicates. For each sample, its assignation coefficient to each of the clusters is represented with a stacked bar chart. The assignation coefficients are represented for analyses ranging from K=2 to 15, the analysis for K=9 (*) is selected as reference. **B.** K optimization procedure based on the entropy statistic for sNMF (left) and the cross-validation error (CV) for Admixture (right). **C.** Breakdown of samples according to genomic dataset. **D.** Breakdown of samples according to sampling location.

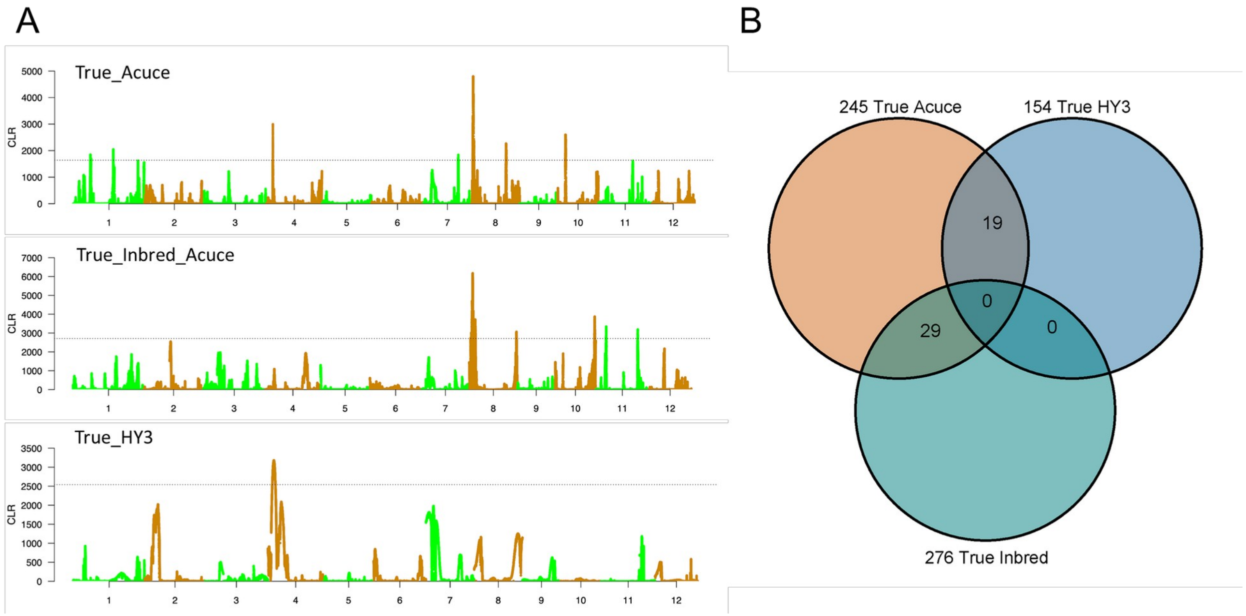


**Fig. S5. Separate breeding histories between Acuce and HY3.**

The lines defined as true Acuce, true Inbred and true HY3 were used to estimate selection signatures genome-wide using SweeD. **A.** CLR scores along the rice chromosomes in the different sub-groups. **B.** Venn diagram of loci under purifying selection (see Table S4 for details).

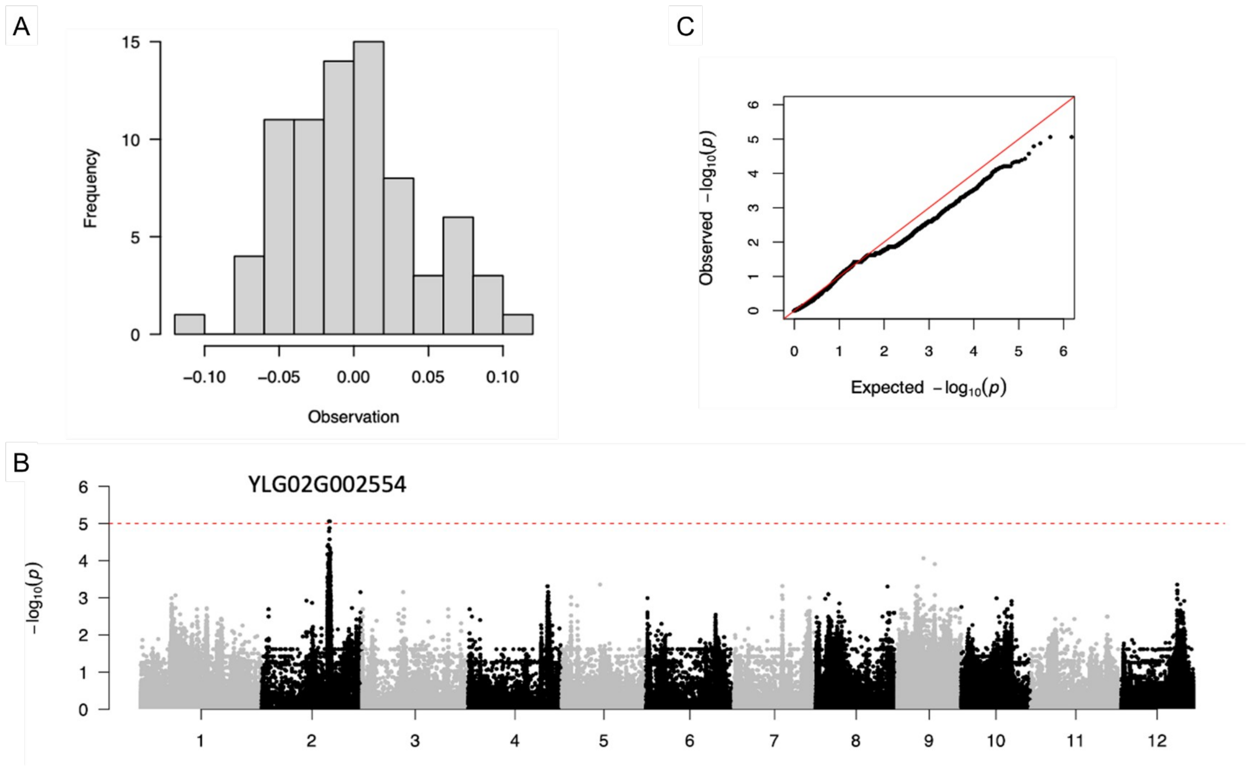


**Fig. S6. GWAS analysis of brown spot resistance in the Acuce landrace.**

**A.** Distribution of the brown spot resistance trait. **B.** Manhattan plot of GWAS for brown spot resistance. The significance threshold is set at -log10(p)=5.0. **C.** Quantile-quantile (Q-Q) plot of GWAS for brown spot resistance.

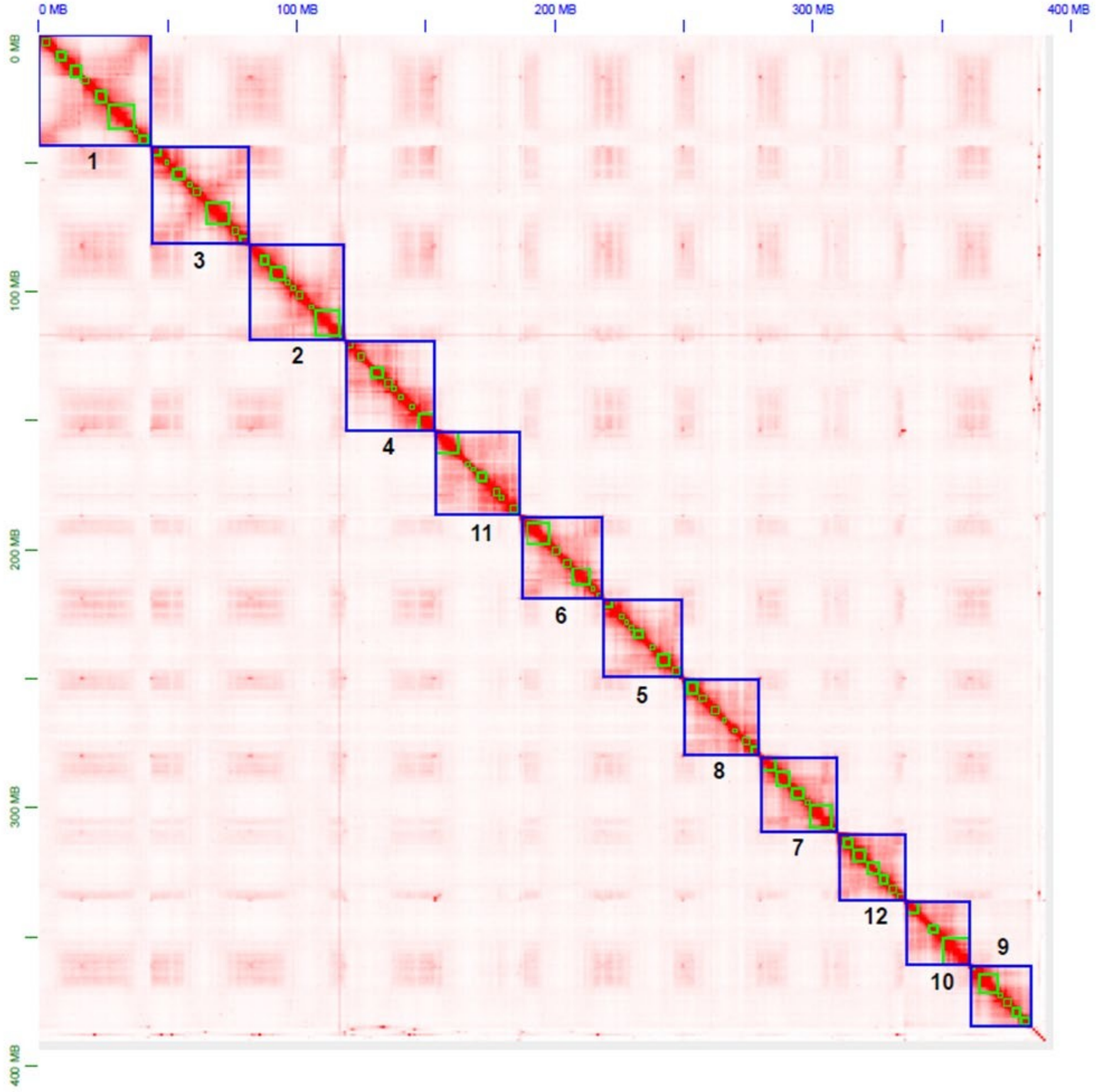


**Fig. S7. Heatmap of Hi-C interactions for Acuce pseudo-chromosomes.**

Numbers correspond to the chromosome numbers used in rice.

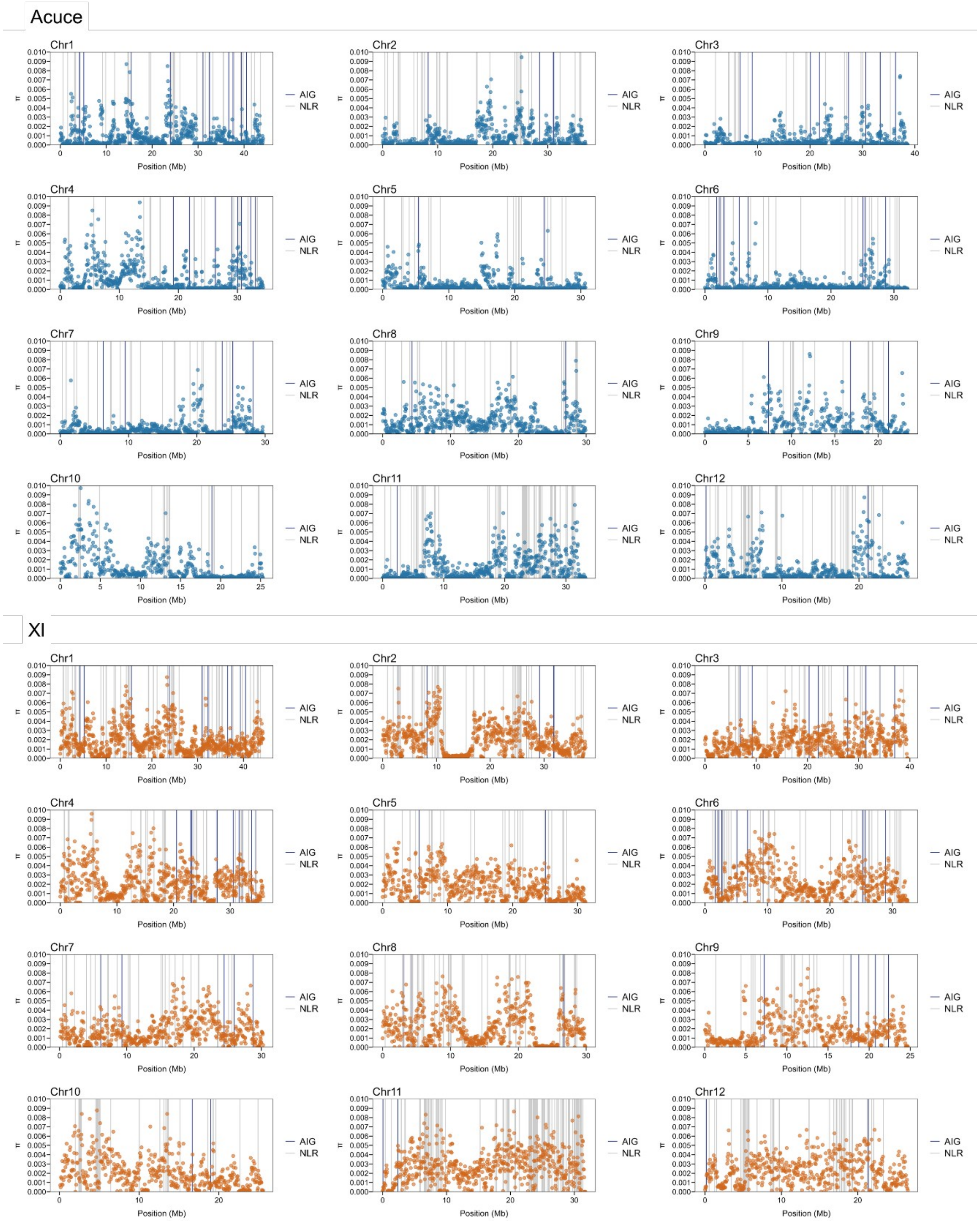
Acuce
XI
Chr1
Chr2
Chr3
Chr4
Chr5
Chr6
Chr7
Chr8
Chr9
Chr10
Chr11
Chr12
π
Position (Mb)
AIG
NLR

**Fig. S8. Diversity feature of Acuce and XI genomes.**

The chromosomal location and π value (in 50 kb windows) for NLR and AIG genes in Acuce and XI populations on each chromosome, based on Acuce15 and R498 genomes, respectively.

**Table S1. Origin and associated metadata for all rice lines used in this study.**

YAU: Yunnan Agriculutral University stock center. 3KGP: 3000 Genomes Project (Wang et al., 2018). Field data: panicles showing neck blast symptoms (sick) or healthy ones were collected, together with the associated rice blast isolate. Susceptibility with P. oryzae was measured under controlled conditions. The GY+avrPia isolate was described in Liao et al. (2016). Acuce and HY3 genomes were first assigned to groups and further delineated in sub-groups for the Acuce and HY3 lines.

Wang W, Mauleon R, Hu Z, Chebotarov D, Tai S, Wu Z, et al. Genomic variation in 3,010 diverse accessions of Asian cultivated rice. Nature. 2018;557:43-49.

Liao J, Huang H, Meusnier I, Adreit H, Ducasse A, Bonnot F, et al. Pathogen effectors and plant immunity determine specialization of the blast fungus to rice subspecies. eLife. 2016;5:e19377.

Available as a separate file.

**Table S2. Diversity statistics per population.**

Statistics computed per group, sub-group, group of village or village. nseff: average number of non-missing alleles per site; S: number of SNPs; π: nucleotide diversity; He: average heterozygosity; Ho: expected heterozygosity; Fis: fixation index; D: Tajima's D. Note that π is reported for the whole dataset.

| Population | nseff | S | Pi | He | Ho | Fis | D |
|---|---|---|---|---|---|---|---|
| Acuce | 216 | 411297 | 74741 | 0.182 | 0.043 | 0.765 | 0.262 |
| Acuce_1 | 116 | 305078 | 50799 | 0.167 | 0.032 | 0.807 | -0.382 |
| Acuce_2 | 88 | 320004 | 59273 | 0.185 | 0.092 | 0.502 | -0.224 |
| Acuce_3 | 12 | 154614 | 51291 | 0.332 | 0.027 | 0.92 | 0.009 |
| inbred | 174 | 428465 | 58369 | 0.136 | 0.046 | 0.665 | -0.716 |
| inbred_1 | 150 | 368213 | 41258 | 0.112 | 0.052 | 0.533 | -1.237 |
| inbred_2 | 6 | 127166 | 65527 | 0.515 | 0.062 | 0.88 | 1.151 |
| inbred_3 | 18 | 158394 | 52526 | 0.332 | 0.166 | 0.501 | 0.603 |
| HY3 | 46 | 365233 | 43824 | 0.12 | 0.07 | 0.418 | -1.751 |
| XI | 99.58 | 774811 | 196523 | 0.254 | 0.035 | 0.864 | 1.071 |
| YYT | 377.19 | 653130 | 169030 | 0.259 | 0.014 | 0.944 | 2.125 |
| YYT1 | 219.56 | 552816 | 134271 | 0.243 | 0.023 | 0.904 | 1.445 |
| YYT2 | 45.96 | 85593 | 10312 | 0.12 | 0.045 | 0.628 | -1.743 |
| YYT3 | 71.72 | 357789 | 113328 | 0.317 | 0.013 | 0.958 | 1.888 |
| YYT4 | 39.78 | 170106 | 15597 | 0.092 | 0.034 | 0.63 | -2.299 |
| V1 | 68 | 299216 | 72527 | 0.242 | 0.03 | 0.878 | 0.571 |
| V2 | 88 | 326834 | 67365 | 0.206 | 0.049 | 0.762 | 0.141 |
| V3 | 58 | 297093 | 67691 | 0.228 | 0.098 | 0.571 | 0.197 |
| Qing Kou | 10 | 179599 | 73648 | 0.41 | 0.02 | 0.951 | 0.804 |
| Xiao Shui Jing | 32 | 191744 | 56526 | 0.295 | 0.016 | 0.946 | 0.727 |
| Tu Guo Zhai | 20 | 243693 | 79929 | 0.328 | 0.013 | 0.962 | 0.687 |
| Xiao Zhai | 6 | 121675 | 54550 | 0.448 | 0.557 | -0.243 | 0.154 |
| Da Yu Tang | 88 | 326834 | 67365 | 0.206 | 0.049 | 0.762 | 0.141 |
| Huang Cao Ling | 42 | 273345 | 71617 | 0.262 | 0.073 | 0.723 | 0.477 |
| Hou Zi Zhai | 2 | 55208 | 55208 | 1 | 1 | 0 | NC |
| Ai Chun | 2 | 78814 | 78814 | 1 | 1 | 0 | NC |
| A Zhe Ke | 8 | 96731 | 37638 | 0.389 | 0.414 | -0.063 | 0.049 |
| Pu Gao Xin Zhai | 4 | 98802 | 54735 | 0.554 | 0.659 | -0.19 | 0.165 |

**Table S3. Genes under purifying selection in true Acuce, Indred and HY3 sub-groups.**

Available as a separate file.

**Table S4. Structure analysis of Acuce and HY3 genomes.**

Available as a separate file.

**Table S5. Differentiation statistics between pairs of populations.**

S: number of polymorphic sites; n1/n2: number of alleles in the two populations; nseff: average number of non-missing alleles per site; Fst: Weir and Cockerham's differentiation statistic; Dj: Jost's D index of differentiation; Gst': Hedrick's differentiation statistic; Da: net pairwise genetic distance; Dxy: raw pairwise genetic distance.

| Pair of populations | S | n1 | n2 | nseff | Fst | Dj | Gst' | Da | Dxy |
|---|---|---|---|---|---|---|---|---|---|
| Acuce_1/Acuce_2 | 386660 | 116 | 88 | 204 | 0.369 | 0.115 | 0.175 | 0.084 | 0.226 |
| Acuce_1/Acuce_3 | 346967 | 116 | 12 | 128 | 0.553 | 0.226 | 0.305 | 0.189 | 0.337 |
| Acuce_2/Acuce_3 | 357302 | 88 | 12 | 100 | 0.452 | 0.18 | 0.261 | 0.141 | 0.296 |
| YYT1/YYT2 | 583061 | 220 | 46 | 265.54 | 0.531 | 0.26 | 0.352 | 0.226 | 0.35 |
| YYT1/YYT3 | 611783 | 220 | 72 | 291.31 | 0.344 | 0.149 | 0.236 | 0.113 | 0.316 |
| YYT1/YYT4 | 581593 | 220 | 40 | 259.44 | 0.469 | 0.22 | 0.312 | 0.183 | 0.311 |
| YYT2/YYT3 | 449913 | 46 | 72 | 117.7 | 0.68 | 0.397 | 0.504 | 0.355 | 0.493 |
| YYT2/YYT4 | 323604 | 46 | 40 | 85.8 | 0.936 | 0.598 | 0.626 | 0.581 | 0.622 |
| YYT3/YYT4 | 474633 | 72 | 40 | 111.6 | 0.68 | 0.407 | 0.513 | 0.362 | 0.498 |
| V1/V2 | 380754 | 68 | 88 | 156 | 0.028 | 0.011 | 0.034 | 0.007 | 0.191 |
| V1/V3 | 340277 | 68 | 58 | 126 | 0.088 | 0.038 | 0.072 | 0.023 | 0.229 |
| V2/V3 | 388852 | 88 | 58 | 146 | 0.184 | 0.063 | 0.111 | 0.041 | 0.215 |
| Xiao Shui Jing/Da Yu Tang | 340304 | 32 | 88 | 120 | 0.007 | 0.007 | 0.034 | 0.005 | 0.187 |
| Xiao Shui Jing/Huang Cao Ling | 296719 | 32 | 42 | 74 | 0.18 | 0.083 | 0.153 | 0.055 | 0.271 |
| Da Yu Tang/Huang Cao Ling | 380597 | 88 | 42 | 130 | 0.158 | 0.056 | 0.105 | 0.037 | 0.219 |
| XI-1/XI-2 | 687215 | 20 | 20 | 39.88 | 0.21 | 0.11 | 0.212 | 0.08 | 0.333 |
| XI-1/XI-3A | 682313 | 20 | 20 | 39.74 | 0.242 | 0.127 | 0.231 | 0.094 | 0.346 |
| XI-1/XI-3B | 638043 | 20 | 20 | 39.91 | 0.267 | 0.14 | 0.249 | 0.103 | 0.351 |
| XI-1/XI-3C | 646648 | 20 | 20 | 39.88 | 0.188 | 0.103 | 0.2 | 0.074 | 0.333 |
| XI-2/XI-3A | 613278 | 20 | 20 | 39.7 | 0.199 | 0.112 | 0.216 | 0.082 | 0.343 |
| XI-2/XI-3B | 559786 | 20 | 20 | 39.88 | 0.209 | 0.117 | 0.223 | 0.085 | 0.347 |
| XI-2/XI-3C | 579827 | 20 | 20 | 39.86 | 0.143 | 0.087 | 0.182 | 0.061 | 0.33 |
| XI-3A/XI-3B | 532930 | 20 | 20 | 39.73 | 0.207 | 0.12 | 0.221 | 0.088 | 0.36 |
| XI-3A/XI-3C | 569777 | 20 | 20 | 39.71 | 0.163 | 0.098 | 0.195 | 0.07 | 0.342 |
| XI-3B/XI-3C | 509263 | 20 | 20 | 39.89 | 0.137 | 0.084 | 0.176 | 0.06 | 0.336 |

**Table S6. Genes under balancing selection in true Acuce, Inbred and HY3 sub-groups.**

For each sub-group (Acuce, Inbred Acuce and HY3), the SNPs under balancing selection are reported. Since several SNPs can be located into one single locus, non-redundant (nr) loci under blanacing selection were extracted for each sub-group. Finally, all loci under balacing selection were concatened into one single table (overlap). AIG: agronomicaly important gene; NLR: nucleotide-binding leucine-rich repeats.

Available as a separate file.

**Table S7. Genome annotation of the Acuce line 15, including NLR genes.**

Available as a separate file.

**Table S8. Nucleotide diversity at AGI and NLR genes.**

Available as a separate file.

**Table S9. Agronomically important genes (AIG) in Acuce genome.**

Available as a separate file.

**Table S10. Presence/absence polymorphism for AIG and NLR genes in Acuce and HY3.**

Available as a separate file.

**Table S11. Genome statistics.**

Available as a separate file.

**Table S12. Statistics of the Acuce genome assembly.**

| Total length (bp) | Total number (bp) | Average length (bp) | N50 (bp) |
|---|---|---|---|
| 46,836,195,431 | 5,609,873 | 8,348 | 13,264 |

**Table S13. Nucleotide diversity per window.**

Available as a separate file.
